# Scaled Testosterone: A Novel Metric to Calibrate Serum Testosterone and SHBG in Men

**DOI:** 10.64898/2026.05.23.727352

**Authors:** DJ Handelsman, GA Wittert, BB Yeap, CM Muir, L Flicker, M Ng Tang Fui, M Grossmann

## Abstract

**Objectives:** Low serum testosterone (T) in men with obesity suggesting T deficiency may be misinterpreted by confounding changes in serum SHBG, T’s circulating carrier protein. Measuring or calculating “free” testosterone (FT) concentrations to define a low T is problematic as cFT is not a valid analytical variable lacking certified standard, quality control or reference range. We developed a novel metric, Scaled Testosterone (ST), comparing standardized serum T (LCMS) and SHBG without invoking hypothetical serum T fractions.

**Methods:** Serum T and SHBG in men (n=10,027) pooled from three population-based studies in Australia were expressed as standardized (Z) scores (ZT, ZSHBG) and their difference ST = ZT-ZSHBG. ST was evaluated in a clinical trial of 51 men with severe obesity undergoing 1 year of diet-induced weight loss.

**Results:** ZT and ZSHBG displayed linear correlation (r=0.44, 10^-11^) with ST approximating zero (-0.33 ±2.14 SD). In non-obese men with low serum T suggestive of organic hypogonadism displayed very low ST indicating ST can evaluate whether a low serum T is proportionate to a concomitant serum SHBG.

In men with obesity, low pre-treatment serum T and SHBG both increased during diet-induced weight loss with no change in serum LH while ST which remained within standard limits at each time.

**Conclusions:** The low serum T in men with obesity may better be considered as the pseudo-hypogonadism of obesity comprising low serum T with proportionately low serum SHBG in the presence of normal serum LH ± FSH serving as a tissue androgen sensor.

## Introduction

Men with obesity typically have a low serum testosterone (T) which may be misinterpreted as T deficiency when viewed in isolation. However, reduced serum T may be due to, and confounded by, changes in serum SHBG, serum T’s carrier protein. Misinterpretation of low T in men with obesity is major contributor to the ongoing global excess of unjustified off-label testosterone prescribing [1], mainly for men without pathologic (structural or genetic) hypogonadism. A common approach to interpret the low T in men with obesity has been measuring or calculating “free” testosterone (FT) [2]. This is based on the based on the longstanding but unproven free testosterone hypothesis which states that unbound testosterone is the bioactive moiety of circulating T; yet unbound T would be equally accessible to site of T degradation so the net effect of whether unbound T is the most or least bioactive moiety is unpredictable [3].Furthermore, FT is not a valid analytical variable as evades the first principle of analytical chemistry -to compare like with like – as FT lacks a certified standard, quality control and valid reference range [4].

Given the theoretical and practical limitations of cFT, a new approach is needed to evaluate low serum T concentrations to determine if they are simply the consequence of a lowered serum SHBG, a prominent feature of male obesity, and to distinguish functional from pathological causes of a low serum T. Therefore, this study aimed to develop an alternative metric, Scaled Testosterone (ST), to compare serum T and SHBG without invoking hypothetical serum T fractions. Using three large scale population-based cohort studies of Australian men to provide both serum T, all measured by liquid chromatography-mass spectrometry (LCMS) in a single laboratory, and SHBG, the standardized (Z) scores for serum T (ZT) and SHBG (ZSHBG) were derived to allow a scaled comparisons of serum T (LCMS) and SHBG embodied in the difference parameter ST (=ZT-ZSHBG).

## Materials and Methods

The assumption-free Scaled Testosterone (ST) metric compares standardized serum T (ZT) and SHBG (ZSHBG) concentrations derived from the population reference data [5] provided by pooling three population-based studies of Australian men [5-7] comprising samples from HIMS study (n=4225), MAILES (n=4531) and CHAMP studies (n=2535) each of which had human ethics approval from their host institutions. Data was pooled from the MAILES study (31/8/2023) and from the HIMS study (15/6/2011) with CHAMP data which originated in the Andrology Laboratory, ANZAC Research Institute. For the present analysis, fasting samples with both serum T and SHBG concentrations and with serum T>5.0 nmol/L excluding only those with likely primary hypogonadism (serum FSH>15 IU/L) comprising samples from HIMS study (n=3911), MAILES study (n= 3535) and CHAMP study (n=2167) samples. Subsets of samples of men with severe obesity (BMI> 35kg/m^2^, n=2145) or with serum T concentrations <5 nmol/L (n=416) or >35 nmol/L, likely to represent pathological hypogonadism or exogenous T administration, respectively, were analysed separately.

To evaluate the ST approach in men with obesity, we used data from the placebo arm (n-51) of an RCT of testosterone vs placebo treatment added to diet in which men with obesity were treated with a hypocaloric diet for 10 weeks followed by 46 weeks maintenance diet [8].Fasting blood sample were taken with serum stored at –80°C until assay using a single LCMS method in one laboratory for all samples reported herein. Serum T was measured by ultrahigh pressure liquid chromatography–mass spectrometry (LCMS) [9, 10] and calibrated against a certified T reference standard (National Measurement Institute, North Ryde, Australia) with a limits of quantification (lowest detectable measurement with CV <20%) of 0.025 ng/ml. Between-run coefficients of variation at four levels (very low, low, medium, and high) of quality control (QC) specimens included were 1.9%–4.5%, 3.8%–7.6%, 2.9%–13.6%, and 5.7%–8.7%, respectively, including all samples from this study. QC samples were run at multiple levels of every run with each new QC control run multiple times in overlap for calibration before use and there was no evidence of assay drift [11, 12]. Serum SHBG, LH and FSH were measured in automated immunoassays (Siemens Immulite for HIMS and MAILES studies, Roche for CHAMP study) all subject to ongoing external quality control (QC) programs with between-assay CV for multiple levels of QC specimens in each run of <6.7% for SHBG, <6.4% for LH and <3.1% for FSH.

## Data analysis

The distributions of serum T and SHBG were evaluated by the Box-Cox transformation approach [13]. The correlation of T with SHBG was evaluated initially as a first-order linear regression and then further evaluation for non-linearity by fitting curvilinear terms (quadratic, cubic) to the linear regression and by non-parametric methods Passing-Bablok regression [14] and LOESS smoothing for graphical depiction [15].

Reference range for the scaled difference (ZT-ZSHBG) was evaluated by the empirical 95% confidence limits of ST. Data was expressed as mean and standard deviation (SD) or median and interquartile range (IQR), as appropriate. All data analysis performed by NCSS 2025 software (NCSS, LLC. Kaysville, Utah, USA).

## Results

Pooled results from the three studies include 10,029 samples having both serum T and SHBG measurements. Subsets evaluated separately included 2145 samples in men with severe obesity (BMI>35kg/m^2^) and additional subsets of 416 men with serum T ≤5.0 nmol/L and 46 men with serum T >35 nmol/L were also analysed separately. This main dataset for analysis included 9613 men with serum T > 5nmol/L comprising 3953 (HIMS study), 3493 (MAILES study) and 2167 (CHAMP study). The 95% confidence limits for the distribution of serum T were 6.6 to 28.4 nmol/L and for serum SHBG 15.4 to 89.7 nmol/L.

For men with serum T > 5.0 nmol, serum T and SHBG had linear correlation (r=0.44, p<10^-11^) across the full range of BMI whether on a natural or log scale (Figure 1). Based on Box-Cox analysis, both serum T and SHBG had close to log-normal distribution and further analyses used the log_10_ derivatives of the analytes in nmol/L units.

**Figure 1.**
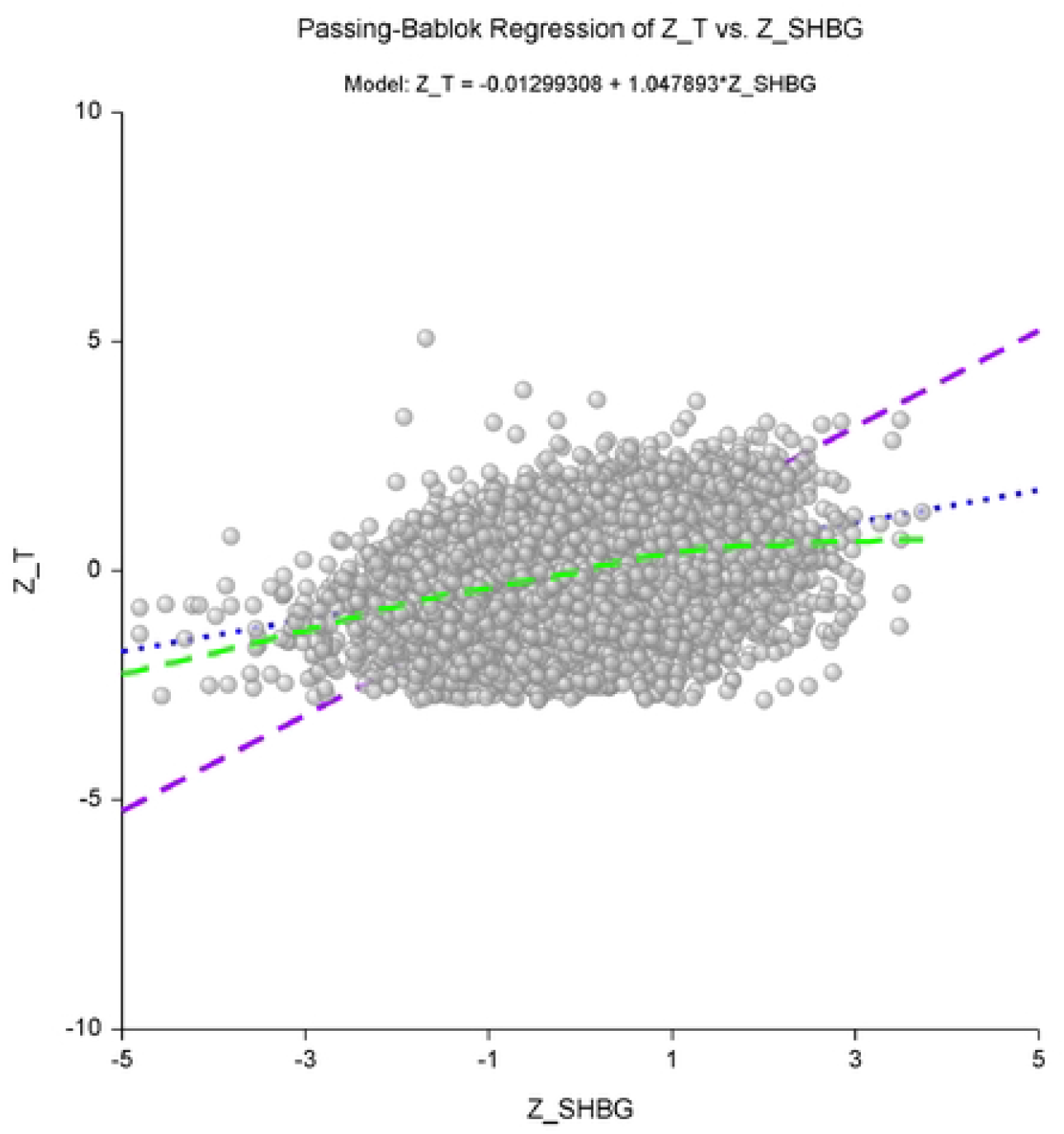
Scatter plot of ZT vs ZSHBG with 9613 samples and linear regression line (dotted line, blue) with the LOESS non-parametric smoothed regression line (dashed line, green) and the non-parametric passing-Bablok regression line (dashed line, purple).

Calculated standardized (Z) scores therefore based on the log-normal distribution were ZT = (log_10_ serum T – 1.15)/0.16

ZSHBG = (log_10_ serum SHBG -1.58)/0.19

From the correlation of ZT and ZSHBG (Figure 1) optimal linear regression parameters were

ZT= -0.013 + 1.048 * ZSHBG with

Intercept = -0.013 (95% confidence limits (CL) -0.014 to -0.010)

Slope = 1.048 (1.019 to 1.078)

In the whole study cohort, ST, the difference between ZT and ZSHBG, was not significantly different from zero (-0.33 ± 2.14 SD, 95% CL -2.7 to 1.80) indicating unbiased agreement overall between ZT and ZSHBG for the reference male population.

Analysis of the hormonal parameters for 9613 men is shown in table 1. Increasing BMI was associated with progressively lower serum T and SHBG, mirrored in their scaled Z scores, ZT and ZSHBG. Notably, ST which remained close to zero with a quantitatively minor upward, rather than downward, trend as BMI increased. Serum LH and FSH displayed quantitatively minor changes with increasing BMI.

**Table 1.**
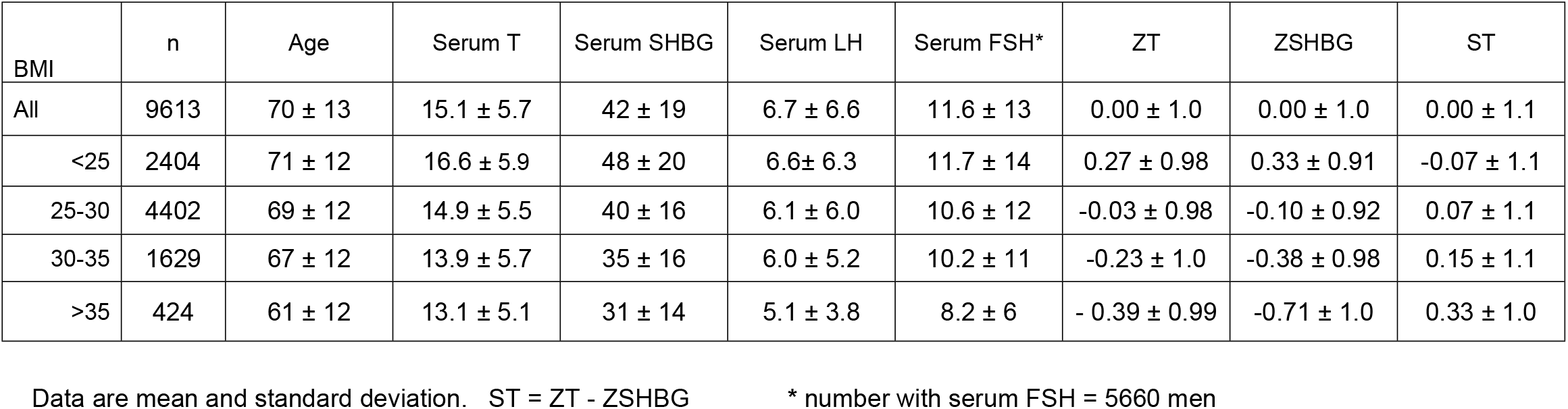
Hormonal characteristics in men with serum testosterone >5 nmol/L.

In the subset of men with obesity (BMI>30 kg/m^2^), serum T and SHBG and their scaled scores decreased with increasing BMI whereas mean serum LH and FSH remained within reference ranges (Table 2).

**Table 2.** Hormonal characteristics in men with obesity (BMI > 30 kg/m^2^)

|  | n | Age | Serum T | Serum | Serum LH | Serum FSH* | ZT | ZSHBG | ST |
| --- | --- | --- | --- | --- | --- | --- | --- | --- | --- |
| All | 2145 | 66 ± 13 | 13.2 ± 6.0 | 34 ± 16 | 5.9 ± 5.0 | 9.6 ± 10 | -0.57 ± 1.9 | -0.46 ± 1.0 | -0.11 ± 2.0 |
| 30-35 | 1695 | 67 ± 12 | 13.4 ± 6.1 | 35 ± 16 | 6.1 ± 5.6 | 10.1 ± 11 | -0.5 ± 1.9 | -0.38 ± 1.0 | -0.15 ± 2.0 |
| >35 | 450 | 62 ± 12 | 12.5 ± 5.6 | 31 ± 14 | 5.1 ± 3.9 | 8.2 ± 6 | -0.7 ± 1.8 | -0.73 ± 1.0 | 0.02 ± 1.9 |

Data are mean and standard deviation. ST = ZT - ZSHBG
\* number with serum FSH = 1524 men

The subset of men (n=416) with low serum T (<5 nmol/L) had a mean BMI 28.3 ± 0.2 kg/m^2^. Their scaled hormonal parameters (ZT, ZSHBG, ST; Table 3) featured very low serum T and negative ZT but with relatively stable minor changes in serum SHBG, ZSHBG and LH so that ST was consistently strongly negative.

**Table 3.** Hormonal characteristics in men with low serum testosterone <5 nmol/L.

| BMI | n | Age | Serum T | Serum SHBG | Serum LH | Serum FSH* | ZT | ZSHBG | ST |
| --- | --- | --- | --- | --- | --- | --- | --- | --- | --- |
| All | 416 | 77 ± 7 | 1.7 ± 1.7 | 45 ± 30 | 9.7 ± 12 | 16.5 ± 20 | -7.9 ± 4.0 | 0.05 ± 1.2 | -8.0 ± 4.0 |
| <25 | 79 | 76 ± 7 | 1.8 ± 1.7 | 52 ± 29 | 9.4 ± 11 | 15.2 ± 16 | -8.7 ± 4.3 | 0.39 ± 1.2 | -9.1 ± 4.7 |
| 25-30 | 191 | 78 ± 6 | 1.5 ± 1.8 | 46 ± 28 | 9.8 ± 12 | 18.9 ± 22 | -7.8 ± 3.9 | 0.06 ± 1.3 | -7.9 ± 4.3 |
| 30-35 | 86 | 76 ± 7 | 1.6 ± 1.6 | 35 ± 16 | 9.0 ± 11 | 7.3 ± 5 | -8.0 ± 3.8 | -0.41 ± 1.1 | -7.5 ± 4.3 |
| >35 | 26 | 72 ± 11 | 2.5 ± 1.7 | 27 ± 12 | 4.6 ± 6 | 8.9 ± 7 | -6.0 ± 3.4 | -1.0 ± 1.0 | -4.9 ± 4.0 |

Data are mean and standard deviation. ST = ZT - ZSHBG \* number with serum FSH = 188 men

A small subset of 46 men with implausibly high serum T (>35 nmol/L) for endogenous T, well beyond the pooled reference limits for serum T by LCMS (7.7 to 29.8 nmol/L) for healthy young men [16], suggestive of exogenous T administration or elevated SHBG for genetic or secondary to liver, thyroid or other disease, where serum T was 42.3 ± 9.4 nmol/L, SHBG was 65 (IQR 48) nmol/L with distinctively high ZT 2.9 ± 0.5 and ST 1.8 ± 1.5 with ZSHBG 1.1 ± 1.2.

Finally, in men with obesity (n=51) in a study of on diet-induced weight loss (placebo arm of an RCT [8]) displayed significant (p<0.001) weight loss with increases of serum T and of serum SHBG during the study by repeated measures linear mixed model analysis with individual data (left panel, Figure 2) and grouped data (right lower panel, Figure 2). The low pre-treatment serum T concentrations (7.0 ± 0.4 nmol/L) increased progressively and significantly (8.6 ± 0.4 at week 10, 10.4 ± 0.4 at week 56) throughout the study. Baseline serum SHBG (23.2 ± 1.5 nmol/L) increased significantly to week 10 (32.4 ± 1.5 nmol/L) but then remained stable at week 56 (29.9 ± 1.6). At all times, serum T and SHBG remained proportionate and within 95% reference limits for ST with no significant divergence (right upper panel, Figure 2). Consistent with the absence of changes in gonadal axis physiology, serum LH concentrations were normal and did not change over the 56 weeks[8].

**Figure 2.**
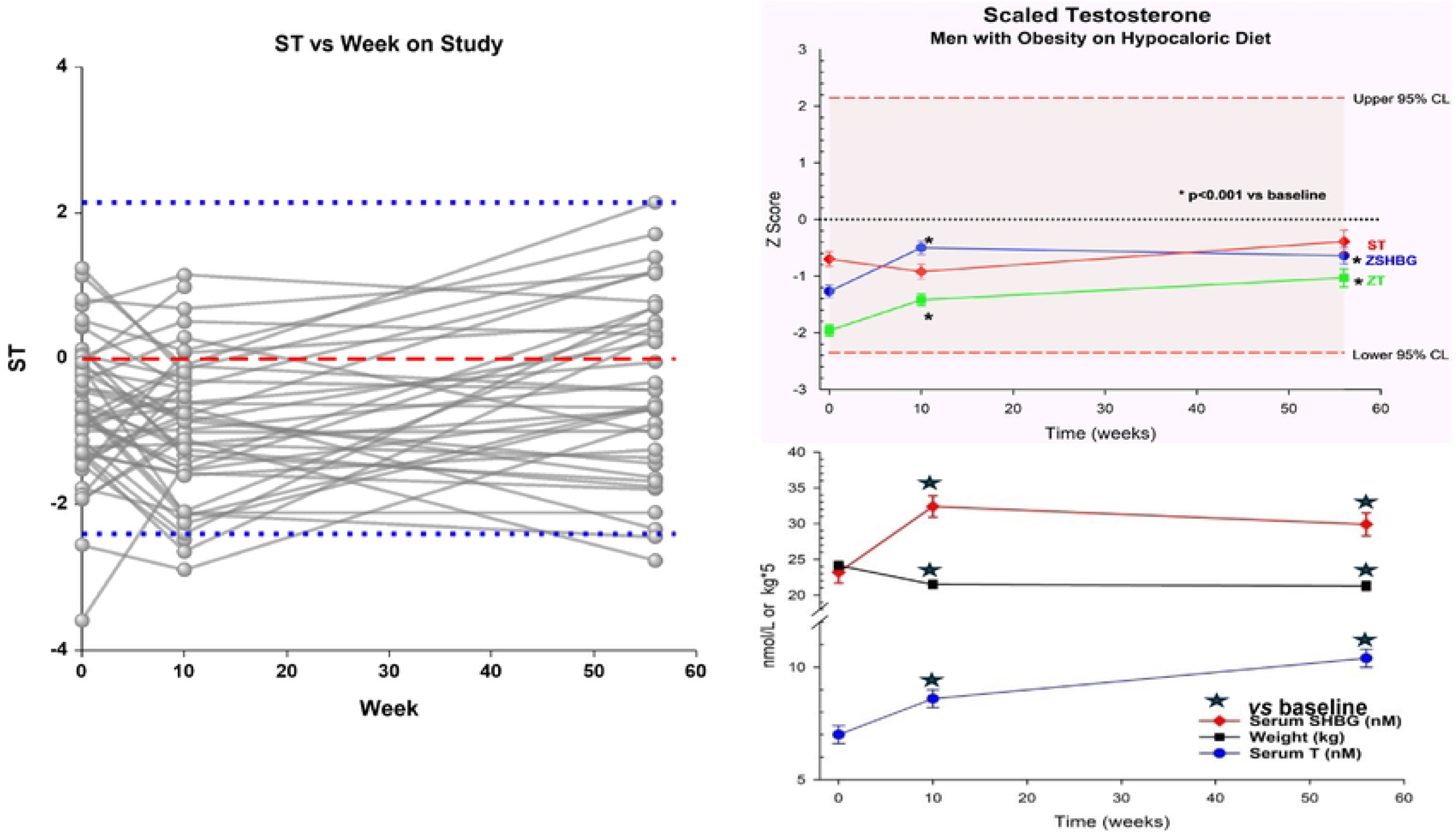
Left panel-ST for 51 individual men with obesity participating in the placebo arm of the 12-month randomized clinical trial of testosterone vs placebo added to hypocaloric diet for 10 weeks followed by a maintenance diet till 56 week. Data is plotted as mean and SEM with 95% confidence limits in blue dotted lines and the zero line of no change in ST in red dashed line. Right panel, upper – Z scores for serum testosterone (ZT, green symbols and line), serum SHBG (ZSHBG, blue symbols and line) and their difference ST = ZT-ZSHBG (red symbols and line). Changes from baseline are significant for ZT and ZSHBG but not for ST during the trial. Right panel, lower – Weight (kg, black symbol and line), serum T (nmol/L, blue symbol and line) and serum SHBG (nmol/L, red symbol and line) against time in weeks on study.

## Discussion

The present study introduces Scaled Testosterone as a novel alternative metric to characterizing the relationship of serum T and SHBG while avoiding invoking hypothetical and unmeasurable fractions of circulating T such as “free” T. The present study, developed at a population scale featuring this new metric at a macro population level, shows its has plausible properties for evaluating androgen status in men with obesity and perhaps other applications where serum SHBG is systematically modified (eg ageing, genetics, liver or thyroid diseases). Further clinical evaluation in smaller, better-defined clinical cohorts of men with and without obesity and SHBG-modifying disease as well as pathological hypogonadism would be required to appraise the clinical utility of this novel metric as an adjunct to evaluating androgen status. This new metric is not introducing a method to measure imaginary fractions of serum T but simply as an aid to determine whether a low serum T is in or out of proportion to that of concomitant serum SHBG. When trying to appraise a low serum T in men with obesity, an appeal to FT is often introduced to determine whether serum T is lowered because of a low serum SHBG or is low regardless of the concomitant serum SHBG. Measuring FT requires laborious manual dialysis methods that are neither robust nor standardized, technically demanding, expensive and rarely available. Recent improvements in dialysis methodology by measuring dialysate T by liquid chromatography-mass spectrometry (LCMS) still provide no quality control over the non-standardized dialysis process itself [17]. These difficulties lead to the usual simpler calculational device of calculating rather than measuring FT (cFT) from serum T and SHBG. However, whether by measured or calculated, FT lacks any certified standard, quality control or valid reference range, thereby failing a key principle of analytical chemistry, that like should be compared with like.

In practice, FT is mostly calculated by various formulae based on either equilibrium binding theory equations [18-20] or assumption-free empirical methods [21, 22]. Equilibrium binding equations involve key assumptions about that the binding affinity of T for SHBG, for which empirical estimates of a single binding site vary over a five-fold range [23] whereas there are two binding sites with different affinities per SHBG molecule [24]. These and other mistaken assumptions influence cFT results leading to discrepancies from assumption-free empirical methods [21]. The theoretical and empirical flaws of FT, including its incoherence in making implausible assumptions that unbound T is more rather than less bioactive, are reviewed elsewhere [3].

Despite these limitations. cFT has been employed in large observational studies attempting to define late-onset hypogonadism [25, 26]) which, among other limitations [27], has had limited utility in that goal due to lack of a certified standard leading to an inability to extrapolate or pool results between labs. Additionally, cFT based on either an empirical or an equilibrium-binding theory formula, failed to predict 24 adverse health outcomes better than measured T by LCMS in a large prospective cohort [12]. Most recently, cFT was not included in recent large placebo-controlled interventional trials using more accurate LCMS T measurement, notably the TTrials [28], TRAVERSE [29] and T4DM [30]; nor is cFT acceptable to justify a Therapeutic Use Exemption for use of testosterone, a prohibited drug in elite sports, for male hypogonadism [31].

These limitations in appraising a low T are particularly salient for men with obesity who usually have a low serum testosterone with proportionate reduction in serum SHBG, T’s circulating carrier protein, but without overt pathologic hypogonadism. In these men a low serum T viewed in isolation may erroneously suggest hypogonadism, a clinical scenario that is among the major drivers of the global excessive testosterone prescribing reported over recent decades [1].

In the present study an alternative metric ST was proposed and developed at a macro (population) scale to evaluate the relationship between serum T and SHBG without reliance on invoking hypothetical and unmeasurable fractions of circulating T. It does not create any derivative of serum T concentration but just whether that circulating concentration is proportionate to serum SHBG or not when proportionate lowering of serum T with normal serum LH and FSH indicates a eugonadal state.

The ST metric demonstrates suitable properties in that there is no bias or divergence between scaled T and SHBG in the general male population, reflecting the overall proportionality of these two scaled measures. It also displays the expected properties that for men with very low testosterone, presumably due to pathological hypogonadism, with scaled T reflecting the disproportionately low circulating T with minimal changes in serum SHBG or serum LH ± FSH (considered as a tissue androgen sensor). This may not be true for the most extreme obesity (BMI>>35 kg/m^2^) which includes men with genetic causes of very severe obesity which may have a hypothalamic pathology (eg Prader-Willi syndrome, genetic insulin resistance)[32-34]. Conversely, in this study the very small population (0.5%) of men with implausibly high serum T (>35 nmol/L) suggestive of either exogenous T administration or a genetic SHBG variant [35], feature high ZT and ST. Furthermore, ST indicates serum T is proportionate to serum SHBG across a wide range of BMI and it is plausible that this reliably distinguish between authentic hypogonadal and pseudo-hypogonadal states. Nevertheless, further studies using smaller, well-defined clinical cohorts of with and without obesity and authentic hypogonadism and SHBG-modifying disease states are required for further validation of the utility of the ST metric.

Among men with simple obesity, ST remained stable while serum LH ± FSH concentrations, operating as androgen sensors, remained within the eugonadal reference range. Hence, a normal serum LH with normal ST may plausibly distinguish non-pathologic pseudohypogonadism due to obesity from pathological hypogonadism and therefore prove useful to dissuade unjustified testosterone treatment in this setting where viewing a low serum T in isolation can create an illusion of a T deficiency state.

Using a large community-dwelling cohort assembled from three Australian population-based studies previously reported with all testosterone measured by LCMS in a single laboratory [5], the present analysis extends to include serum SHBG. It also considers as separate subsets men with obesity (BMI>30 kg/m^2^) and those with low serum T (< 5.0 nmol/L) suggestive of pathological hypogonadism. The latter restriction is plausible compared with the best estimate of the lower limits of the reference range for healthy young men using LCMS measurements without pathologic hypogonadism of about 7.5 nmol/L [16], based on the pooled estimates of available studies. The prevalence of serum T<5 nmol/L (416/10,029, 4%) is higher in this older population than the estimated prevalence of pathologic hypogonadism in the general male population (0.5%)[36], presumably due to the older age group acquiring more co-morbidities which creates the progressive reduction in serum T well known to be associated with male ageing [12].

The present novel approach builds on the concept that serum T is reduced in obesity due to reduced circulating SHBG. The mechanism of reduced hepatic SHBG production in obesity remains unexplained although multiple potential contributions have been suggested. These include hepatic insulin resistance possibly involving increased circulating monosaccharides [37] and/or reduced adiponectin stimulation of HNF-4α [38].

An important consequence of the present findings is that evaluation of a low serum T associated with obesity or otherwise should include measurement of serum SHBG in the same sample. This is necessary to elucidate the significance of the low serum T but any diagnosis of hypogonadism requires further clinical and laboratory investigations, notably serum LH and FSH, to determine if there is a cause of pathologic hypogonadism warranting testosterone treatment. These findings also support a better description of the low serum T in men with obesity as the pseudo-hypogonadism of obesity comprising mildly low serum T with proportionately low serum SHBG and normal serum LH ± FSH, serving as tissue androgen sensors [27, 39]. It is not clear how well this ST approach would work in men with genetic severe obesity syndromes [40] which may have an additional hypothalamic component so that gonadotropins may not be reliable indicators of androgen exposure. It should be emphasized that ST does not represent any measure of circulating T comparable with cFT with the aim of defining androgen deficiency, but only whether the circulating T level is proportionate to circulating SHBG.

Limitations of this present study include that this evaluation of the ST concept at macro population scale (without individual clinical details on the participants) rather than at a clinical scale. Hence further studies to evaluate this new metric at a clinical scale, including comparison with cFT, are needed to distinguish between functional causes of low serum T due to lowered serum SHBG and pathological hypogonadism.

## Acknowledgements

Nil

ANZAC Research Institute Concord Hospital, NSW 2139 Australia

E

## Notes

### Competing Interest Statement

I have read the journal's policy and the authors of this manuscript have the following competing interests: DJH is an Academic Editor for PLOS One but played no role in the evaluation of this manuscript. The other authors have no relevant disclosures.

